# Filament dynamics driven by ATP hydrolysis modulates membrane binding of the bacterial actin MreB

**DOI:** 10.1101/2021.04.08.439044

**Authors:** Vani Pande, Nivedita Mitra, Saket Rahul Bagde, Ramanujam Srinivasan, Pananghat Gayathri

**Author notes:** Corresponding author: Pananghat Gayathri.

## Abstract

MreB, the bacterial ancestor of eukaryotic actin, is responsible for shape in most rod-shaped bacteria. While the eukaryotic actin utilizes ATP hydrolysis to drive filament treadmilling, the relevance of nucleotide-driven polymerization dynamics for MreB function is unclear. Here, we report mechanistic insights into the interplay between nucleotide-binding, ATP hydrolysis and membrane-binding of *Spiroplasma citri* MreB5 (ScMreB5). Antiparallel double protofilament assembly of ScMreB5^WT^ with ATP, ADP or AMPPNP and an ATPase deficient mutant ScMreB5^E134A^ demonstrate that the filaments assemble independent of ATP hydrolysis. However, capture of the filament dynamics revealed that efficient filament formation, bundling through lateral interactions and filament disassembly are affected in ScMreB5^E134A^. Hence, the catalytic glutamate (Glu134 in ScMreB5) plays a dual role – it functions as a switch by sensing the ATP-bound state for filament assembly, and by assisting hydrolysis for triggering disassembly. Glu134 mutation also exhibits an allosteric effect on membrane binding, as observed from the reduced liposome binding compared to that of the wild type. Thus, ATP hydrolysis can modulate filament length and bundling, and consequently the orientation of MreB filaments on the cell membrane depending on the curvature. Binding of ScMreB5 with liposomes is mediated by surface charge-based interactions, demonstrating paralog and organism specific features for MreB function. We conclude that the conserved ATP-dependent polymerization and disassembly upon ATP hydrolysis has been repurposed for modulating curvature-dependent organization of filaments on the membrane.

## Introduction

The chromosomally encoded bacterial actin, MreB plays a pivotal role in cell shape determination in bacteria (1, 2). Genetic studies have shown that the deletion of *mreB* genes leads to loss of rod shape and eventual lysis of non-spherical bacteria (3–5). It functions as a scaffold for the assembly of cell wall synthesis machinery and, thus, locally leads to cell wall insertion favoring a rod shape (6–8). During growth, the short filaments of MreB align approximately perpendicular to the long axis of the rod-shaped cells (7, 9). Their circumferential movement recruits the peptidoglycan synthesis machinery at uniformly distributed locations along the long axis, thus reinforcing the rod shape.

A characteristic feature of MreB filaments is their ability to bind to the lipid bilayer or monolayer *in vitro* (10–12). Hence, MreB filaments are capable of sensing as well as generating membrane curvature in liposomes, independent of the peptidoglycan synthesis machinery (1, 11, 13). Experimental studies on *Escherichia coli* (EcMreB), *Caulobacter crescentus* (CcMreB) and *Thermatoga maritima* (TmMreB) MreBs have shown that MreB interacts with the cell membrane either via N-terminal amphipathic helix in gram-negative bacteria and/or a hydrophobic loop in subdomain 1A in gram-positive bacteria (11).

MreB filaments possess an antiparallel double protofilament assembly (8, 10), as opposed to the parallel protofilament arrangement in most actin family members such as eukaryotic actin (14) and ParM (15), an actin-like protein in plasmid segregation. Biochemical studies of TmMreB have shown that it is an active ATPase (16). Light scattering studies for MreBs have shown that it polymerizes in the presence of ATP, AMP-PNP (non-hydrolyzable analog of ATP), GTP or ADP (16–19). Additionally, *Bacillus subtilis* MreBs (BsMreB) were shown to undergo nucleotide independent polymerization (18). Further, double protofilament assembly was observed in the crystal structure of CcMreB without nucleotide (10). Therefore, the significance of nucleotide binding or hydrolysis for filament formation and dynamics for MreB function is ambiguous.

Though *in vivo* effects of ATP hydrolysis mutants of MreB have indicated a potential role for hydrolysis in MreB function (20–22), the exact role of hydrolysis-dependent filament dynamics is unknown. Mutational defects in the ATP binding pocket of MreB altered the localization of MreB filaments, cell morphology and chromosome segregation in *B. subtilis* and *C. crescentus* (21, 23). Further, the spatial regulation of MreB filaments in response to cellular curvature has been hypothesized to depend on hydrolytic activity (2, 24). The observation of filament dynamics through an *in vitro* reconstitution approach has not been reported for MreB, probably due to the challenges with imaging the short filaments using light microscopy.

Until recently, studies on MreB function, including cell shape regulation and maintenance, cell division and motility, were reported only from cell-walled bacteria (3, 21, 23), wherein it functions in conjunction with the peptidoglycan synthesis machinery. Therefore, how rod-shape is achieved in bacteria without a cell-wall remains enigmatic. Interestingly, many of the features of MreBs from cell-walled bacteria, such as the antiparallel double protofilament assembly and membrane-binding are common to MreB from a wall-less bacterium too, as demonstrated by our study on MreB5, one of the 5 paralogs of MreB in a wall-less helical bacterium *Spiroplasma citri* (25). Role of multiple paralogs (5 to 7) of MreBs within these organisms remains poorly understood. Recent work from our lab showed that among the 5 paralogs of MreB, MreB5 (ScMreB5) is essential for the helical shape and motility of *S. citri* (25). Orientation of MreB filaments within cells has been proposed to be dependent on the cell diameter (8). The narrow diameter of *Spiroplasma* cells (∼100 – 150 nm) (26) compared to most other bacterial species makes the investigation of *Spiroplasma* MreB paralogs especially interesting. These aspects prompted us to carry out an in-depth study of ScMreB5. Characterization of an MreB, which functions independent of cell wall synthesis machinery in a bacterium of unusually thin diameter, will help in identifying the fundamental mechanism and conserved signatures for MreB function.

Here, we report the structural and biochemical characterization of ScMreB5^WT^ (wild type). An ATPase-deficient mutant of ScMreB5, ScMreB5^E134A^, shows defects in polymerization compared to ScMreB5^WT^. Assisting in conformational changes during polymerization is an additional novel role proposed for the catalytic residue, which has always been implicated only in stimulating hydrolysis in most actin family members such as actin (14), ParM (15) and MamK (27). Furthermore, through lipid specificity studies and mutational analysis, we show that the positively charged C-terminal residues play a major role in facilitating electrostatic interactions for liposome binding. These observations provide novel insights into the role of ATP hydrolysis and its effect on conformational dynamics and membrane binding properties, which are essential for MreB function. The results also highlight the conserved features of allostery and filament dynamics observed in both MreB and actin, despite the differences in their protofilament organization.

## Results

### Crystal structure of ADP-bound ScMreB5 shows a conserved single protofilament organization

We recently reported the crystal structure of ScMreB5 bound to AMP-PNP (PDB ID: 7BVY; (25)). ScMreB5 also crystallized in the presence of ADP (PDB ID 7BVZ). The overall structure of ADP-bound ScMreB5 (ScMreB5–ADP) was very similar to AMP-PNP bound ScMreB5 (ScMreB5–AMP-PNP; superimposing with an RMSD of 0.87 Å; Fig. 1A-B). Clear electron density was observed for the nucleotides in both ADP and AMP-PNP bound states (Fig. 1C-D, data collection statistics in Table S1). The packing of ScMreB5–ADP molecules in the crystal structures revealed a single protofilament assembly, similar to that of ScMreB5–AMP-PNP (Fig. 1E; (25)). The longitudinal repeat distance of 51.1 Å for both these protofilaments were remarkably same when compared with TmMreB (28) and CcMreB (10) subunit repeat distances of 51.1 Å, despite all the four belonging to different space groups and packing environments. The hydrophobic loop in the IA domain, predicted to be the membrane-binding loop in TmMreB (11), was disordered (residues 93 – 97) in ScMreB5–ADP (Fig. 1A). This loop was ordered in the ScMreB5–AMP-PNP structure (Fig. 1B), probably due to crystal packing differences. The domain-wise comparison of the ScMreB5 structures with CcMreB is shown in Table S2.

**Figure 1.**
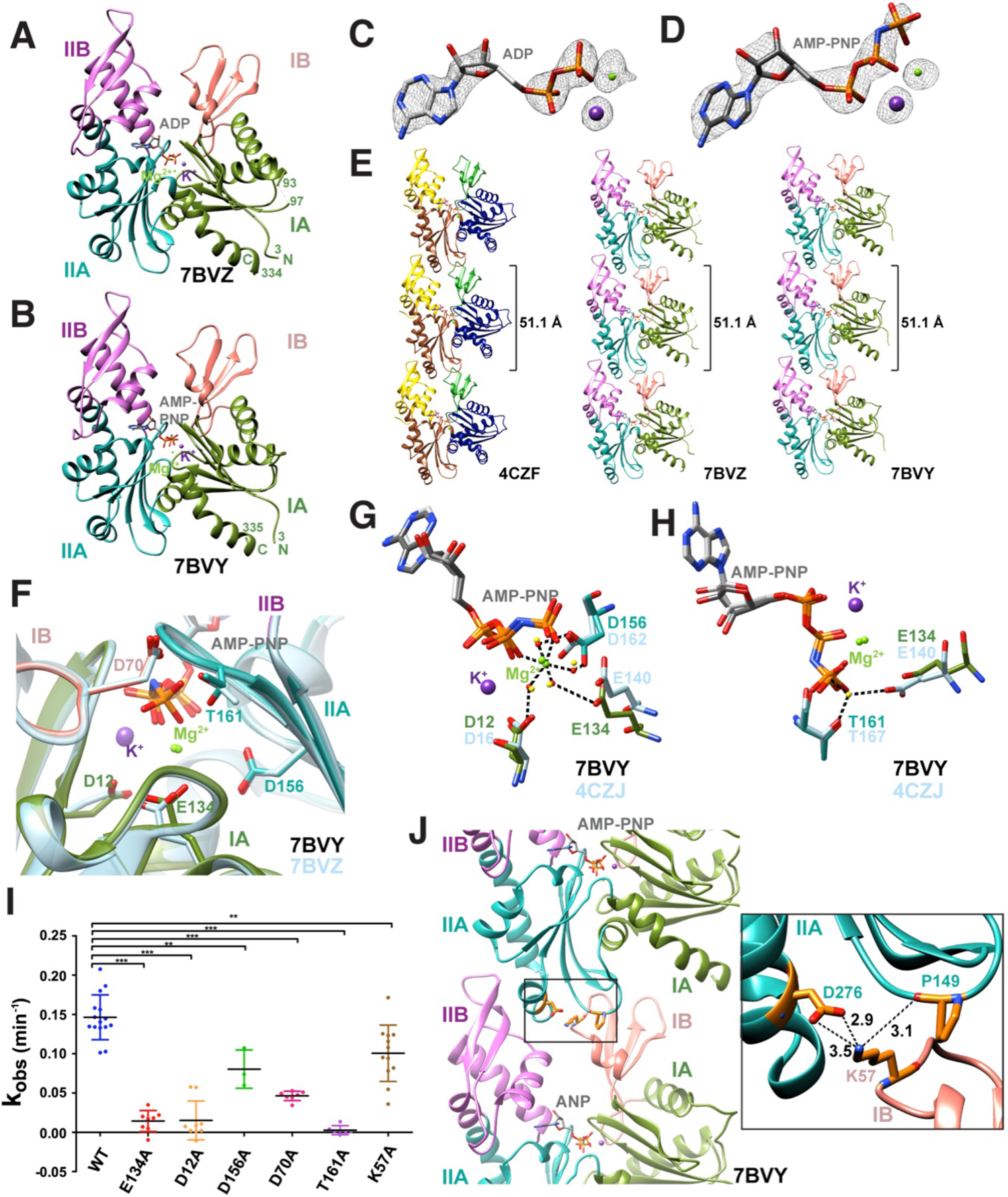
ScMreB5 possesses a conserved protofilament arrangement and is an active ATPase. A, B. Crystal structures of ScMreB5 in ADP (PDB ID: 7BVZ) and AMP-PNP bound states (PDB ID: 7BVY). The sub-domains IA, IB, IIA and IIB are colored and labeled. N- and C-terminal ends are labeled ‘N’ and ‘C’ respectively and the terminal residue numbers marked. The chain breaks in 7BVZ are also labeled by their residue numbers (93 and 97). **C, D.** Electron density for the bound ADP and AMP-PNP with Mg^2+^ and K^+^ (composite omit map F_o_ –F_c_ shown at 2.0 σ). **E.** Protofilament structures of CcMreB (PDB ID: 4CZF), ScMreB5 with bound ADP (PDB ID: 7BVZ) and AMP-PNP (PDB ID: 7BVY). Both the nucleotide bound structures of ScMreB5 have the same subunit repeat as CcMreB (51.1 Å) in their protofilament assemblies. **F.** Zoomed view of the residues at the nucleotide binding pocket. Residues of ScMreB5–AMP-PNP (PDB ID 7BVY; domain-wise colours) are shown superimposed with corresponding residues in ScMreB5–ADP (PDB ID 7BVZ; blue-grey). **G.** Residues involved in Mg^2+^ coordination in ScMreB5 (Asp156, Glu134 and Asp12). Distances for Mg^2+^ coordination are marked by dotted lines for ScMreB5–AMPPNP. **H.** Residues adjacent to the γ-phosphate, Glu134 and Thr161, at the nucleotide binding pocket. Distances with the catalytic water are marked by dotted lines for CcMreB structures. **I.** ATPase activity characterization of ScMreB5. *k_obs_* (min^-1^) for the ScMreB5^WT^, active site mutants and the polymerization mutant [ScMreB5^WT^ (WT; N=3; n=15), ScMreB5^E134A^ (E134A; N=3; n=8), ScMreB5^D12A^ (D12A; N= 2; n=7), ScMreB5^D156A^ (D156A; N=2; n=3), ScMreB5^D70A^ (D70A; N=2; n=7), ScMreB5^T161A^ (T161A; N=1; n=8) and ScMreB5^K57A^ (K57A; N=2; n=12)]. The error bar denotes mean with standard error of the mean (unpaired t-test, two tailed, *** denotes p<0.0001; ** denotes p = 0.0011-0.0019). 10 µM protein, 1 mM ATP and 1 mM MgCl_2_ were used in this assay. N denotes the number of independent protein purification batches, while n denotes the total number of repeats. **J.** Intra-protofilament polymerization interface of ScMreB5 (PDB ID: 7BVY). Inset: A zoomed in view of the interface showing the residue Lys57 of subdomain IB interacting with Asp276 and Pro149 of subdomain IIA. The distances in Å are labeled for the interactions.

An interesting observation in both the structures of ScMreB5 (ADP and AMP-PNP complexes) was the presence of strong electron density for a potassium ion, positioned between α and β phosphates of ADP and AMP-PNP (Fig. 1A-D), providing a structural basis for increased stability of ScMreB5 in KCl buffer (Fig. S1A-E). The crystal structure of CcMreB has a water at the position equivalent to the potassium ion binding site in ScMreB5 (Fig. S1F). The residues Asp12 and Asn17, which are involved in potassium ion coordination in ScMreB, are well conserved among MreBs from different bacteria (Fig. S2) and bind to the corresponding water molecule in CcMreB.

### ScMreB5 is an active ATPase

Residues in the nucleotide binding cleft of ScMreB5 are well conserved with respect to other MreBs and yeast actin (Fig. 1F; Fig. S2). In ScMreB5, the water molecules that form the coordination sphere for Mg^2+^ are held together by the side chain carboxyl oxygens (OD1/OD2 or OE1/OE2) of Asp12, Asp156 and Glu134 (Fig. 1G). Electron density for the catalytic water, typically situated at an in-line geometry with the γ-phosphate moiety, was absent in the structure of ScMreB5–AMP-PNP, and hence not modelled. However, Glu134 and/or Thr161 might interact with the catalytic water, a hypothesis based on structure alignments with other MreB and actin structures in their double protofilament conformations (Fig. 1H; Fig. S3 A-B) (10, 29, 30).

We carried out ATPase activity measurements of ScMreB5^WT^ and active site residue mutants, ScMreB5^D12A^, ScMreB5^E134A^ and ScMreB5^D156A^, by measuring the released phosphate using a colorimetric assay (Fig. 1I). We also chose to mutate Asp70 (ScMreB5^D70A^; Fig. 1F), which is well conserved in MreBs whereas in actin it is replaced by His73 (Fig. S3C) (31). Studies on actin have shown that His73 is important for polymerization and for regulating the phosphate release upon ATP hydrolysis (32). All the ScMreB5 mutants studied showed a decrease in ATPase activity, compared to the wild type (Fig. 1I). *k_obs_* values for ScMreB5^WT^ and the mutants are tabulated in Table S3. ScMreB5^WT^ exhibited similar activity as shown earlier for EcMreB and TmMreB (17, 33).

Crystal structures of the various states of CcMreB showed that conformational changes upon polymerization affects the positioning of the catalytic residues at the active site (10). Hence, in addition to mutants of the ATP-binding pocket residues, we checked the ATPase activity of a polymerization interface mutant ScMreB5^K57A^. Lys57 is present at the intra-protofilament interface of ScMreB5 protofilament assembly (Fig. 1J). This residue is conserved across different MreBs (Fig. S2). ATPase activity of ScMreB5^K57A^ was less compared to ScMreB5^WT^ (Fig. 1I; Table S3), indicating an allosteric communication between polymerization interface and the active site.

### ScMreB5 **^E134A^** filaments exhibit impaired dynamics

We attempted to study the nucleotide-dependence of filament formation for ScMreB5^WT^ and its ATP hydrolysis mutant ScMreB5^E134A^ by observing the presence of filaments *in vitro* using cryo-electron microscopy (cryo-EM). ScMreB5^WT^ in the presence of ATP (Fig. 2A) and AMP-PNP (Fig. 2B) formed high density of double protofilament assemblies having a sheet-like appearance of laterally associated filament bundles. We also observed filaments of ScMreB5^WT^ in the presence of ADP (Fig. 2C) and ScMreB5^E134A^ in the presence of AMP-PNP (Fig. 2D). However, very few sheet-like bundles were observed in ScMreB5^WT^–ADP and ScMreB5^E134A^–AMP-PNP compared to ScMreB5^WT^–ATP and AMP-PNP (Fig. 2A-D). While filaments observed for ScMreB5^WT^–ADP and ScMreB5^E134A^–AMP-PNP demonstrated that ATP hydrolysis was not required for filament formation *in vitro*, observation of a very few sheet-like bundles suggested lower filament density or defective bundling or both.

**Figure 2.**
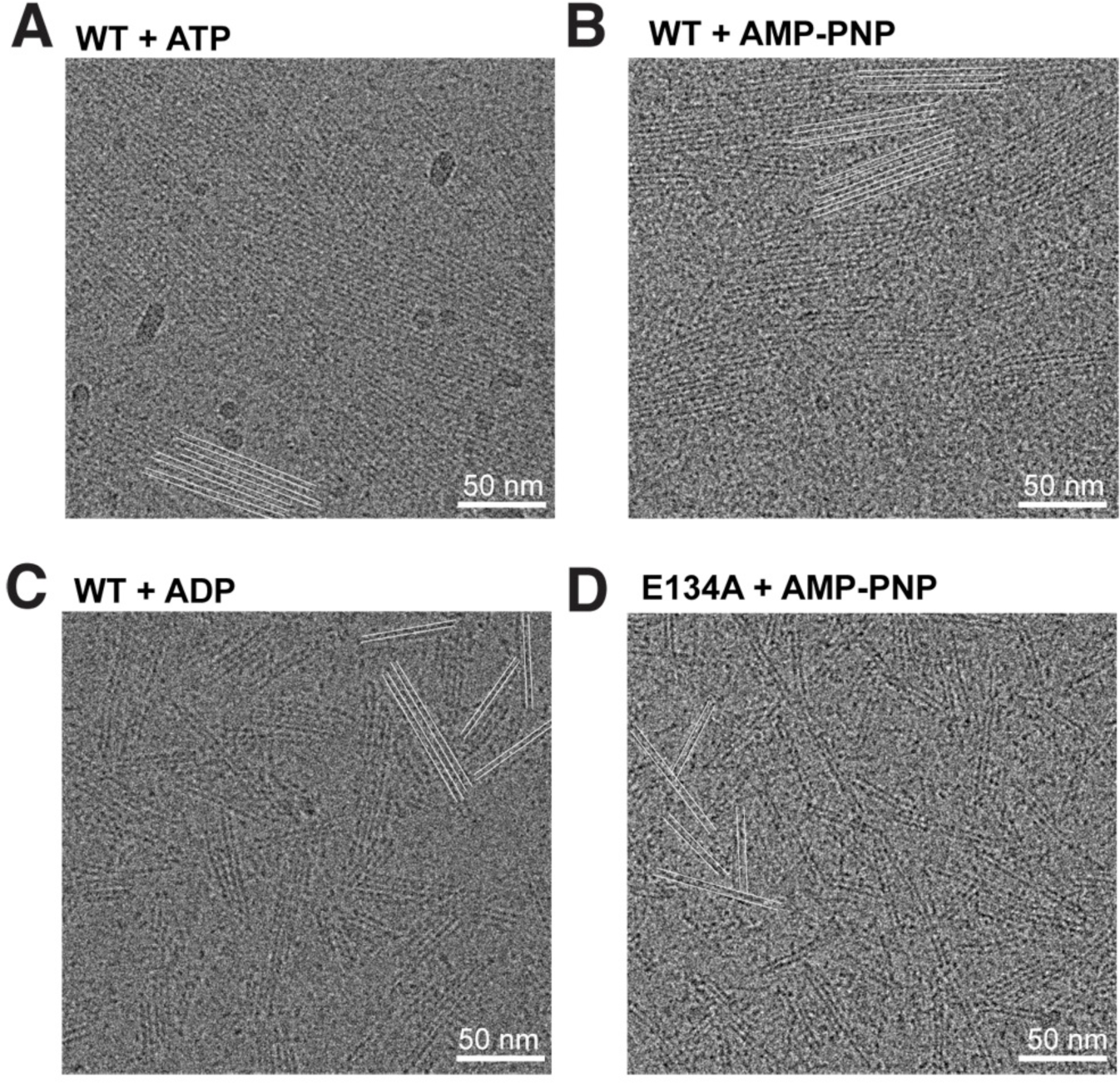
ScMreB5 filaments form double protofilament assemblies independent of nucleotide hydrolysis. Cryo-electron micrographs showing filaments of **A.** ScMreB5^WT^ in the presence of 5 mM ATP and MgCl_2_ **B.** ScMreB5^WT^ in the presence of 5 mM AMP-PNP and MgCl_2_ **C.** ScMreB5^WT^ in the presence of 5 mM ADP and MgCl_2_ **D.** ScMreB5^E134A^ mutant (hydrolysis deficient) in the presence of 5 mM AMP-PNP and MgCl_2_ A few double protofilaments are highlighted by pairs of parallel white lines to enable easy visualization of the filament distribution. Concentration of protein used was 50 µM. Scale bar denotes 50 nm.

With an aim to visualize polymerization dynamics of ScMreB5 filaments, we expressed N-terminal GFP-fusion constructs of ScMreB5^WT^ and ScMreB5^E134A^, respectively, in fission yeast and monitored their filament assembly. We had earlier reported dynamics of EcMreB polymerization in fission yeast with a similar N-terminal GFP-fusion (22). ScMreB5^WT^ showed filaments extending across the cells that would eventually bundle up along the long axis of the cells (Fig. 3A, 3B, Movie S1). However, unlike ScMreB5^WT^ filaments, ScMreB5^E134A^ filaments appeared less dense in yeast cells, and were defective in bundling (Fig. 3A). Defect in bundling of filaments was more clearly visible by super-resolution imaging (3D-SIM) of ScMreB5 filaments (Fig. 3B; Movie S1). Moreover, filaments were observed in very few cells expressing ScMreB5^E134A^ in comparison to ScMreB5^WT^(Fig. 3C).

**Figure 3.**
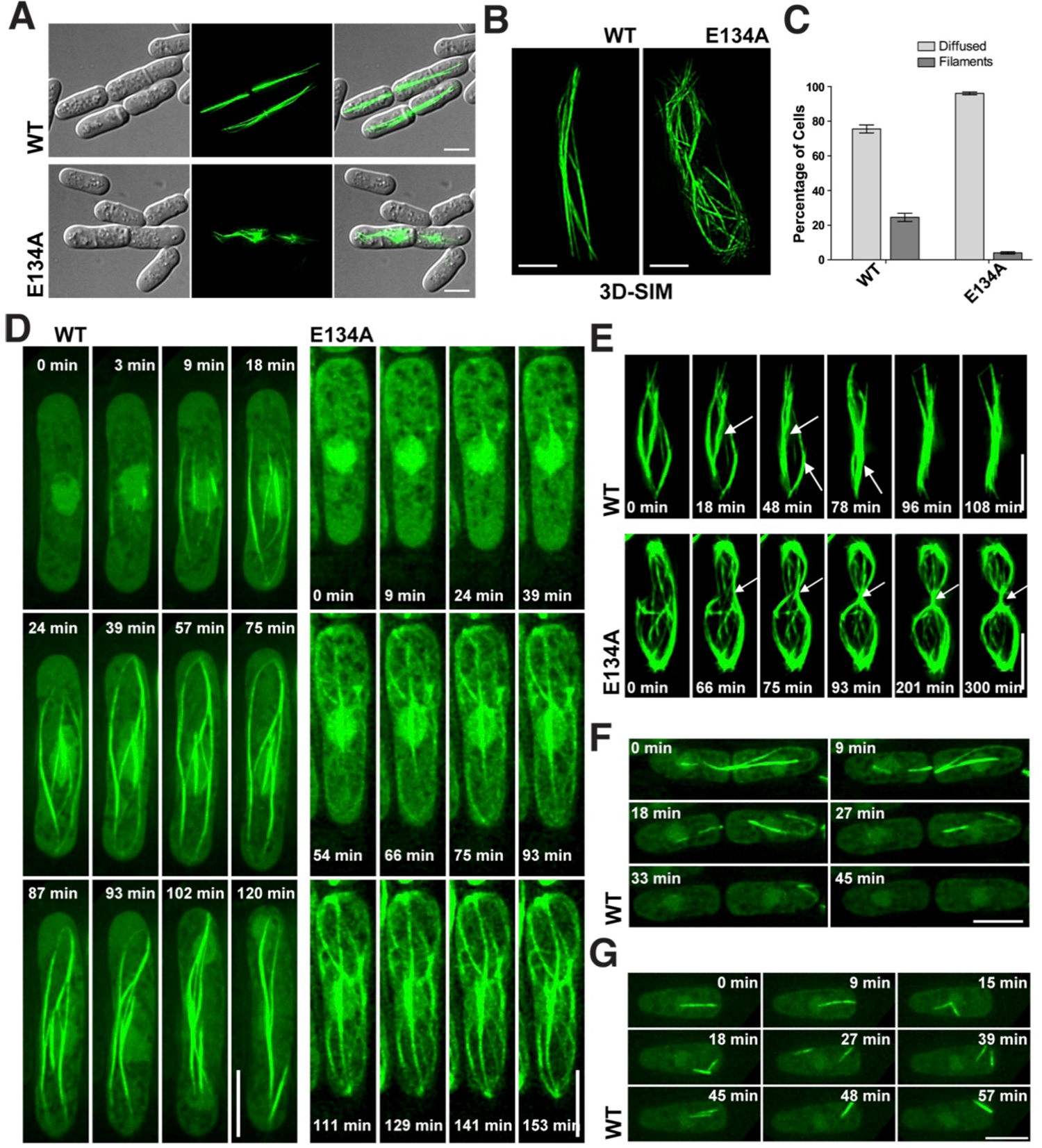
Expression of GFP-tagged ScMreB5 in *S. pombe* cells reveals filament assembly defects of the ATP hydrolysis mutant ScMreB5^E134A^. **A.** ScMreB5^WT^ and ScMreB5^E134A^ with N-terminal tagged GFP expressed in *S. pombe* cells. Representative images are shown for the DIC images of the cell boundaries (left column), a GFP channel (central column) and overlay (right column). **B.** 3D-SIM images of ScMreB5^WT^ and the ScMreB5^E134A^ mutant are shown (related to Movie S1). Filaments of ScMreB5^WT^ appear as bundles spanning from one end to the other end of the cell whereas ScMreB5^E134A^ filaments are not tightly bundled. **C.** Plot comparing the percentage of cells showing either diffuse fluorescence or filaments for ScMreB5^WT^ and ScMreB5^E134A^. Cells were grown in EMM without thiamine for 36 – 48 hours. Cells expressing ScMreB5^E134A^ show more diffused fluorescence and very less filaments in comparison to the ScMreB5^WT^. The mean values (n=3) are plotted and the error bar denotes mean with standard error of the mean. **D.** Time lapse microscopy showing polymerization for ScMreB5^WT^ and ScMreB5^E134A^ (related to Movie S2). **E.** Time lapse microscopy showing bundling for ScMreB5^WT^ and defect in bundling for ScMreB5^E134A^ filaments (related to Movie S3). **F.** Time lapse microscopy of cells expressing ScMreB5^WT^ undergoing division (related to Movie S4). Filament disassembly can be observed post cell division. **G.** Time lapse microscopy of cells expressing ScMreB5^WT^ shows fragmentation and reannealing or bundling of filaments (related to Movie S5). Scale bar denotes 5 µm. For panels **A** – **B** and **D** – **G**, cells were grown in EMM without thiamine for 24 – 32 hours before imaging. Cells were placed on an agarose pad as described in Materials and Methods and imaged at every 3 minute time interval.

Time-lapse imaging of polymerization of ScMreB5^WT^ and ScMreB5^E134A^ within yeast cells confirmed that polymerization and bundling were efficient in ScMreB5^WT^ compared to ScMreB5^E134A^ (Fig 3D; Movie S2). Polymers in cells expressing ScMreB5^E134A^ were observed much later in time compared to those expressing ScMreB5^WT^ (Fig. 3D). This is consistent with the observation that filaments of ScMreB5^E134A^ were seen in fewer cells compared to ScMreB5^WT^ (Fig. 3C), suggesting the requirement of a higher concentration of monomers for polymerization of ScMreB5^E134A^. Bundling of ScMreB5^WT^ was often promoted by cell septation as the ingressing septa brought the filaments in close proximity. The bundling defect of ScMreB5^E134A^ was clearly seen in yeast cells undergoing cell division (Fig. 3E; Movie S3). Interestingly, ScMreB5^WT^ filaments, in many instances, disassembled in the daughter cells immediately after a cytokinesis event (Fig. 3F; Movie S4). In a few cases, filaments of ScMreB5^WT^ appeared to undergo fragmentation and reannealing (Fig. 3G; Movie S5). However, we did not observe any such events of disassembly of the ScMreB5^E134A^ filaments and they appeared stable. Filament stabilization is a characteristic feature of ATPase defective mutants of the actin family. The electron microscopy images and the filament dynamics appeared to suggest decreased polymerization and bundling for ScMreB5^WT^ in the presence of ADP and for ScMreB5^E134A^ irrespective of the nucleotide present.

### Surface charge and active site mutation influences liposome binding of ScMreB5

Recently, we showed that ScMreB5 interacts with liposomes (25). We further explored the sequence determinants and lipid specificities that influence membrane-binding of ScMreB5 in this study. Though ScMreB5 lacks a distinct amphipathic helix at its N- or C-terminal ends (Fig. 4A, 4B), it possesses Ile95 and Trp96 in the hydrophobic loop of the IA domain (Fig. 4C), which might act as membrane anchors (Fig. 4C, 4D) (34). Hence, we made single (ScMreB5^I95A^ and ScMreB5^W96A^) and double mutant (ScMreB5^I95A,W96A^) constructs of these residues of ScMreB5^WT^ and tested their binding using liposomes having lipid composition resembling *S. citri* membrane (35). In comparison to ScMreB5^WT^, the mutations did not abrogate liposome binding significantly. Both the single and double mutant proteins were found in the pellet fraction (Fig. 4E, 4F). This suggested that the hydrophobic loop might not serve as the sole membrane anchor for ScMreB5.

**Figure 4.**
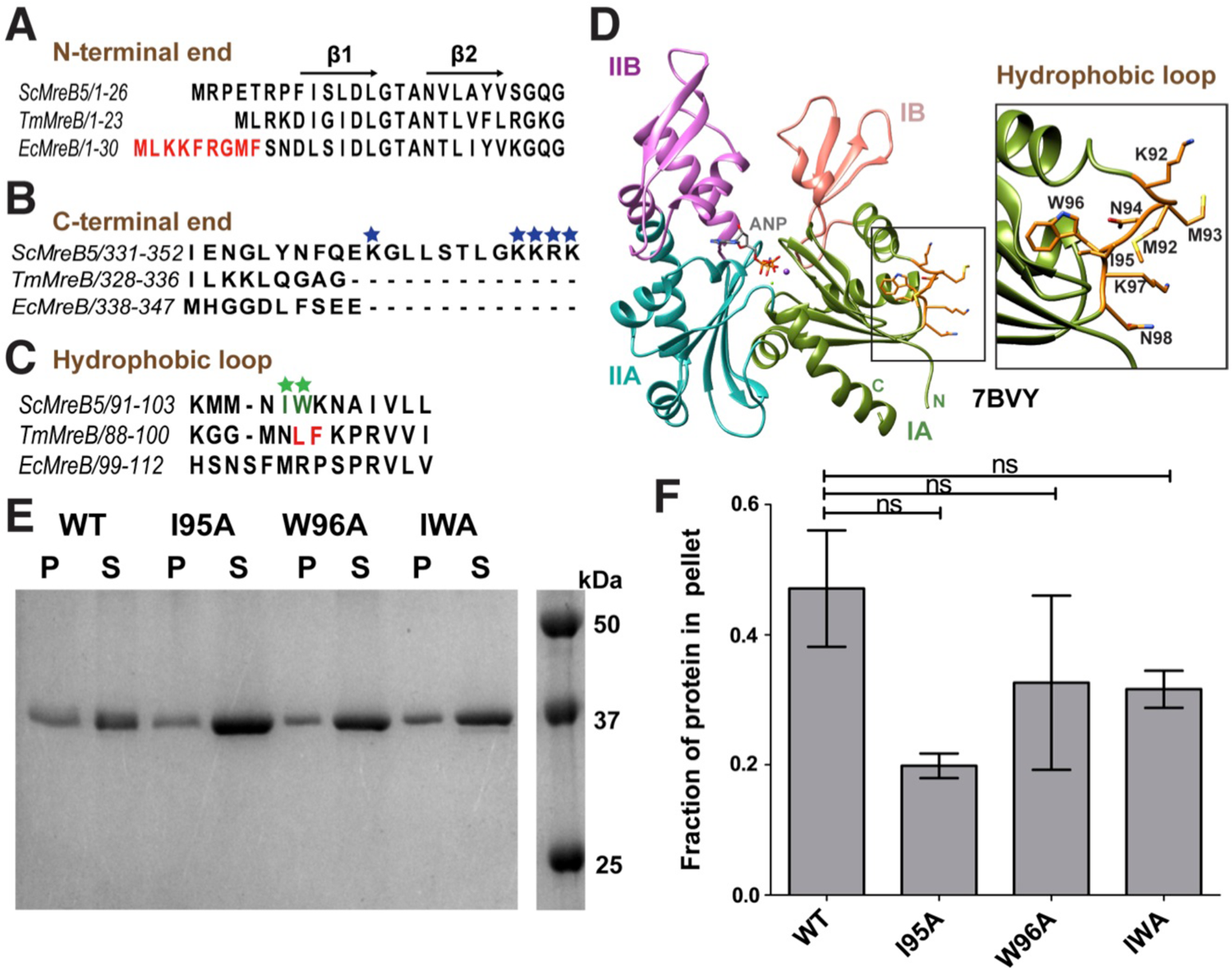
ScMreB5 binds to liposomes. **A.** Multiple sequence alignment of ScMreB5 with *T. maritima* (Tm) and *E. coli* (Ec) MreBs showing the absence of amphipathic helix at the N-terminus. Amphipathic helix of EcMreB is highlighted in red. Secondary structures are labeled on top of the alignment. **B.** Sequence alignment of C-terminal region of ScMreB5 with Tm and Ec MreBs shows longer C-terminal tail enriched with positively charged residues (highlighted with blue stars). **C.** Multiple sequence alignment of ScMreB5 with TmMreB and EcMreB in the region of hydrophobic loop. The residues interacting with the membrane for TmMreB and predicted residues for ScMreB5 are highlighted in red and green (marked with stars), respectively. **D.** Crystal structure of AMPPNP-bound ScMreB5 (PDB ID: 7BVY) with proposed membrane insertion loop (orange) in domain IA (green). Inset: A zoomed view of the loop. The N and C-terminal ends of ScMreB5 are labeled as N and C respectively. **E.** A representative 12% SDS-PAGE gel of liposome pelleting assay for comparing membrane binding of ScMreB5^WT^ and the hydrophobic loop mutants (single mutants ScMreB5^I95A^ and ScMreB5^W96A^ and double mutant ScMreB5^I95A, W96A^ denoted as I95A, W96A and IWA, respectively). P and S represent the pellet and supernatant fractions of the reaction. Concentrations of liposomes and protein used in the assay are 1 mM and 2 µM respectively. **F.** Plot showing relative intensities of the fraction of protein in the pellet corresponding to ScMreB5^WT^ and hydrophobic loop mutants calculated from the SDS-PAGE gels (representative gel shown in panel **E**) from 3 independent experiments. The error bar denotes mean with standard error of the mean (unpaired t-test; ns denotes p>0.05).

In order to decipher mechanistic details of ScMreB5–liposome interaction, we carried out a lipid specificity study. *S. citri* membrane consists of ∼38% phosphatidylglycerol (DOPG, an anionic lipid) and ∼14% phosphatidylcholine (DOPC, a neutral lipid) (35). Hence, we tested whether liposome binding by ScMreB5 could be charge-specific. ScMreB5^WT^ and the hydrophobic loop mutants did not bind to 100% DOPC liposomes (the protein was entirely obtained in the supernatant of the pelleting assay; Fig. 5A), whereas binding was observed for 100% DOPG liposomes (the protein was present in the pellet fraction; Fig. 5A, Fig. S4A). Interestingly, MreB5s in spiroplasmas have a longer C-terminal end, which contains a stretch of lysines and arginines (Fig. 5B). Based on the structures of MreBs, we know that both the N- and C-termini of the protein and the hydrophobic loop face the same side of the monomer and filament surface (Fig. 4D), though the C-terminus is unstructured in the ScMreB5 crystal structures (Fig. 1A). The presence of positively charged residues suggested that a charge-based interaction might be mediated by the C-terminal tail. We tested this hypothesis by a liposome pelleting assay with 100% DOPG liposome for ScMreB5^ΔC10^ (ScMreB5 construct with the last 10 residues deleted), compared to ScMreB5^WT^ (Fig. S4A, B).

**Figure 5.**
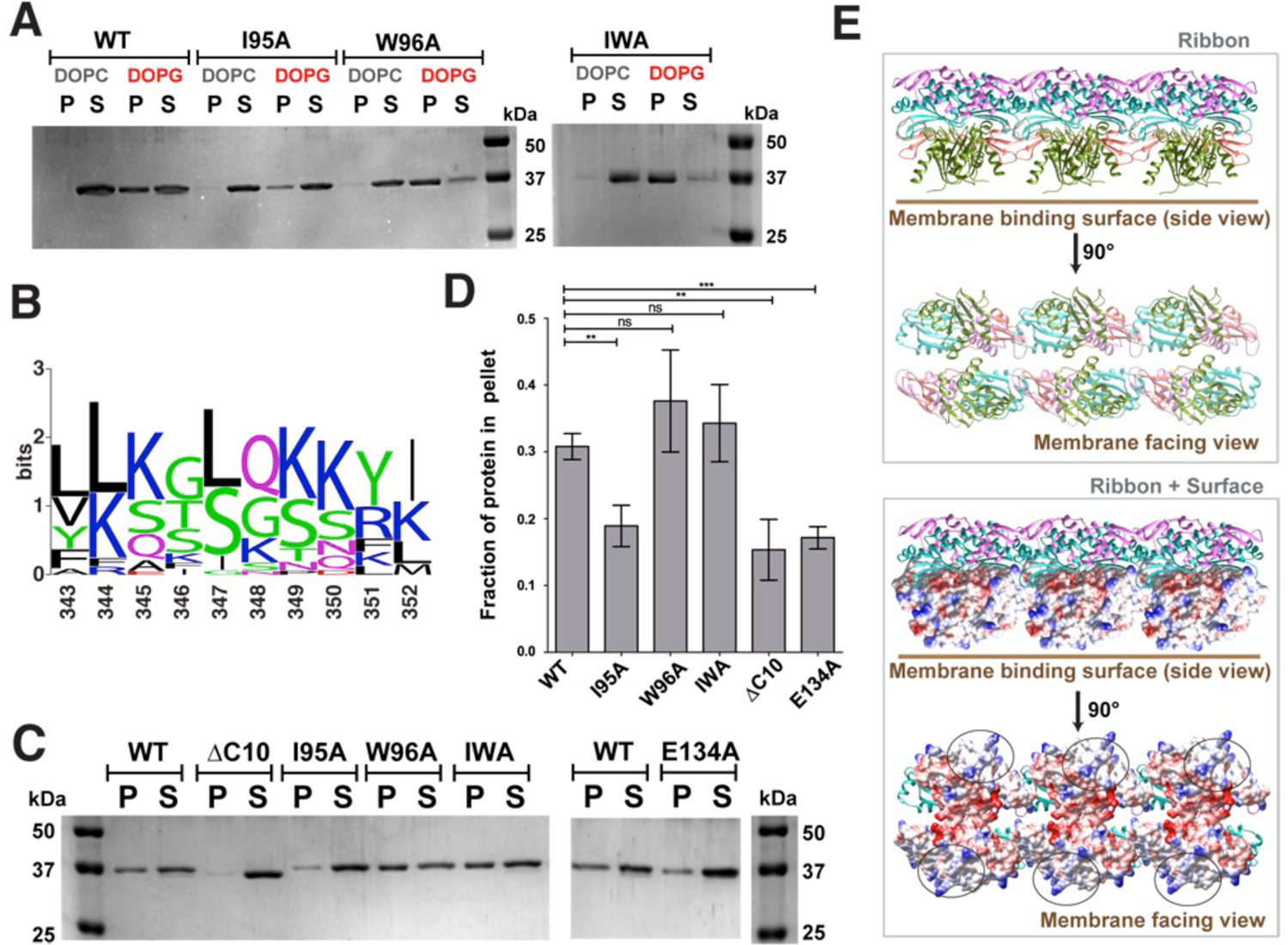
Membrane binding of ScMreB5 is modulated by lipid charge and conformational changes driven by Glu134. **A.** A representative 12% SDS-PAGE gel of liposome pelleting assay for determining membrane binding of ScMreB5^WT^ and the hydrophobic loop mutants with a neutral lipid DOPC and an anionic lipid DOPG. P and S represent the pellet and supernatant fractions of the reaction. Concentrations of DOPG and DOPC liposomes and protein used in the assay are 1 mM and 2 µM respectively. **B.** Weblogo of C-terminal end, showing the presence of lysine and arginine in *Spiroplasma* MreB5s. The numbering on the x-axis is with respect to the last ten residues of ScMreB5. **C.** A representative 12% SDS-PAGE gel of liposome pelleting assay showing a decrease in binding for ScMreB5^WT^ and mutants (ScMreB5^ΔC10^, ScMreB5^I95A^, ScMreB5^W96A^, ScMreB5^I95A, W96A^, ScMreB5^E134A^ denoted as ΔC10, I95A, W96A, IWA and E134A respectively) at 20%:80% (DOPC:DOPG) liposome ratio. P and S represent the pellet and supernatant fractions of the reaction. Concentrations of liposomes and protein used in the assay are 600 µM and 2 µM respectively. **D.** Plot showing relative intensities of the fraction of protein in the pellet calculated from the SDS-PAGE gel from at least 4 independent experiments (representative image in panel **C**). The error bar denotes the mean with standard error of the mean (unpaired t-test, ns denotes p>0.05; *** denotes p<0.0001; ** denotes p ranging from 0.0011 to 0.0019). **E.** Different views of membrane binding face of double protofilament ScMreB5. Electrostatic surface potential of the membrane binding face (IA and IB subdomains; bottom box) of double protofilament of ScMreB5 is shown corresponding to the ribbon views of the double protofilament (top box). Circled regions within the surface show the regions of positive and neutral charge for the membrane-binding face of the filament. The double protofilament of ScMreB5 was modelled using CcMreB double protofilament, PDB ID 4CZE. Subdomains IA and IB are coloured pink and light green.

To comparatively analyse liposome binding by the mutants, we chose a fixed concentration of liposomes based on a binding curve obtained for ScMreB5^WT^ with increasing concentrations of liposomes (Fig. S4C). 600 µM, a concentration just below saturation in the binding curve (Fig. S4C), was maintained as a constant liposome concentration for further assays. Next, we repeated the pelleting assays by varying ratios of DOPC:DOPG in the liposome preparation to tease out the contributing factors to membrane binding (Fig. S4D). Based on this, 80% DOPG at 600 µM of liposomes was used for the pelleting assays (Fig. S4C, D).

ScMreB5^ΔC10^ exhibited reduced binding of the protein to 600 µM liposomes with 80% DOPG (Fig. 5C, D). From the above result, it appeared that membrane binding by ScMreB5 was mainly driven by a positive charge distribution on the membrane-binding surface, and was not restricted to a small stretch of residues. The surface potential of the membrane binding face of the modelled ScMreB5 double-protofilament is also consistent with this hypothesis, which is either hydrophobic or positively charged (Fig. 5E). Next, we explored if the liposome interaction of ScMreB5^E134A^ is affected by the mutation. Interestingly, liposome binding assay of ScMreB5^E134A^ showed a significant decrease compared to that of ScMreB5^WT^ (Fig. 5F-G), although the residue is situated away from the membrane-binding surface. The effect observed was similar to that of other loop mutants at the membrane surface such as ScMreB5^I95A^, thus suggestive of an allosteric communication between the nucleotide binding pocket and membrane-binding interface of MreB.

## Discussion

Among the actin filament family members, MreB filament is unique due to the antiparallel arrangements of the protofilaments, implying the absence of a kinetic or structural polarity between the two ends. Structures of MreB filaments have highlighted that the conformational changes accompanying filament formation is very similar to those observed in actin and ParM. The active site residues, including the residues coordinating Mg^2+^ and those required for optimal orientation of the catalytic water, are highly conserved in MreBs, actins and across the Hsp70 superfamily members. Our ATPase activity measurements point out a role for these residues in ATP hydrolysis, thus emphasizing that ATP hydrolysis is an inevitable feature for MreB (as well as ScMreB5) function. Effect of the polymeric interface mutant ScMreB5^K57A^ and the inter-subdomain contact mutant ScMreB5^D70A^ on the ATPase activity proves the existence of an allosteric communication between the ATP-binding pocket and the polymerization interface in MreB too, a conserved feature of many characterized actin family members.

A thorough analysis of the reported crystal structures of CcMreB (10) showed us that the catalytic glutamate (Glu140 in CcMreB; Glu134 in ScMreB5) functions as an interaction hub, forming a network of interactions with gamma phosphate of the nucleotide, catalytic water and residues from various MreB sub-domains (Fig. 6). The entire network of interactions with all the 4 subdomains was observed only in the double filament conformation and not in the single protofilament or monomeric states (Fig. 6, inset). The gamma phosphate of the nucleotide and Glu140 side chain play key roles in the network. Thus, the residue acts as the sensor for the ATP-bound state and triggers double protofilament conformation – an important feature of nucleotide-state dependent polymerization. Additionally, the requirement of Glu140 in providing an optimal active site geometry for ATP hydrolysis hints that the residue also triggers efficient ATP hydrolysis, thus playing a crucial role in nucleotide-dependent polymerization dynamics through initiating disassembly. Thus, Glu140 drives the conformational switch between the ATP-bound double protofilament state and the ADP state incompatible with the double protofilament conformation. The crystal structures of ScMreB5 capture the single protofilament conformations of Glu134 (corresponding to Glu140 of CcMreB).

**Figure 6:**
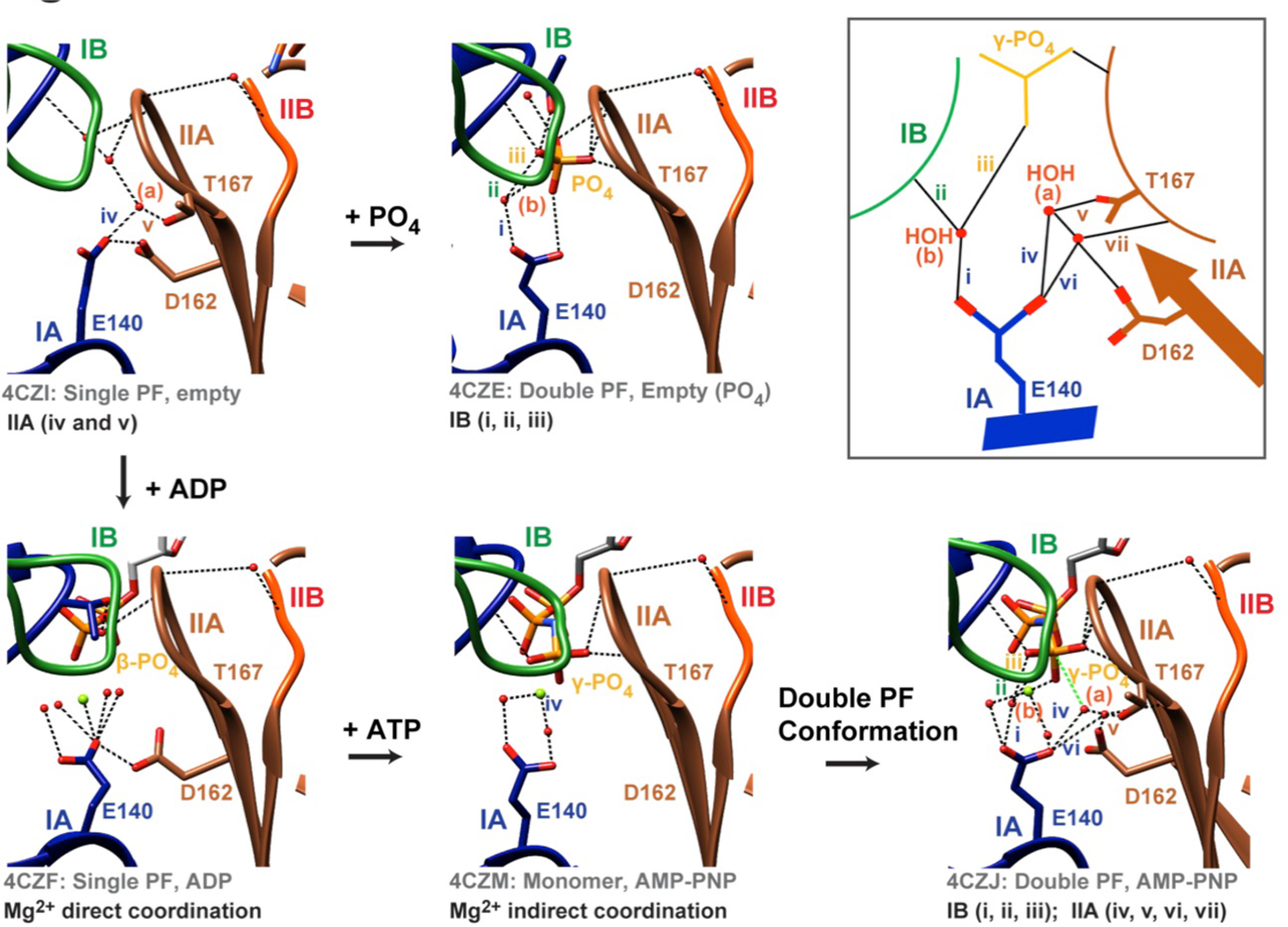
Mechanism of ATP dependence of MreB function Glu140 (CcMreB numbering) plays a pivotal role in ATP-dependent conformational change of MreB. Schematic (box): Glu140 in the double protofilament AMP-PNP bound state (PDB ID 4CZJ) holds the IB and IIA subdomain via water mediated interactions, (a) and (b). These water mediated interactions are not present for ADP single filament (PDB ID 4CZF) and AMPPNP monomeric state (PDB ID 4CZM) where Glu140 functions only in Mg^2+^ coordination.

The antiparallel double protofilament structure of MreB filament highlights a repetitive arrangement within the filament, without a twist angle between monomers, ideally favoring a planar lipid membrane interaction (10, 11). How these filaments align against curved membrane surfaces and/or bring about a curvature in liposomes is enigmatic. While there are theoretical models on how this might be achieved (36), our study based on ScMreB5^E134A^ provides insights into the role of ATP-driven dynamics in polymerization and membrane binding of MreB, a hypothesis earlier put forward based on molecular dynamics simulations (37). Impairment of membrane binding by ScMreB5^E134A^ indicates that the conformational change facilitated by Glu134 on sensing gamma phosphate of ATP is required for efficient liposome interaction, possibly mediated by the filaments bundling through lateral interactions. Sheets of filaments with lateral interactions might also be helpful in promoting binding and modulating the membrane curvature.

ScMreB5 is a major cytoskeletal protein that confers helical shape and facilitates motility in *S. citri* (25). Hence, the mechanistic basis of ScMreB5 function is of special interest to understand the minimal functional requirements of MreBs in bacterial cell shape determination. In ScMreB5, we report a novel mode of membrane binding of MreBs assisted by surface charge-based interactions. Earlier studies on MreBs (*T. maritima* and *E. coli* MreBs) identified the membrane interaction to be abrogated completely by the mutation of the hydrophobic loop or the N-terminal amphipathic helix (11). Our results show that this mode of interaction is not universal for MreBs. Our liposome binding studies with ScMreB5 indicate that charge-mediated interactions contribute to the membrane binding features of ScMreB5. The lipid chemistry and the surface properties of ScMreB5 highlight organism-specific or paralog-specific modes of membrane binding based on sequence variations within a conserved filament architecture. These features can potentially modulate the energetics of filament interaction with the membrane, thereby affecting orientation of MreB filaments necessary for modulation of cell diameter and shape. In the absence of membrane attachments facilitated by the peptidoglycan synthesis related proteins such as RodZ (38), a surface extensive interaction with the membrane might help in orienting the filaments in a cell-wall-less organism.

We demonstrate the allosteric effect of ATP binding and hydrolysis residues on efficient filament formation and membrane binding. Our results emphasize that ATP hydrolysis is essential for a conformational switch leading to filament dissociation and disassembly. Our observations highlight that the fundamental features of MreB are conserved, independent of the involvement of the peptidoglycan synthesis machinery. We envisage a mechanism in which bundles of ATP-bound MreB filaments sense an optimal curvature for binding, remodel the membrane and hydrolyze ATP, and then exchange ADP with ATP and bind to the adjacent region with favourable curvature for binding, thus resulting in a processive motion. *Spiroplasma* MreB5 provides an excellent prototype not only for understanding the conserved features across all MreBs, but also for understanding the role of ATP hydrolysis in curvature sensing, helicity generation and motility driven by the filaments. Further studies of ScMreB5 in *S. citri* will ascertain the mechanistic basis of the role of ATP hydrolysis in motility and helicity.

## Materials and Methods

### Cloning

*Spiroplasma citri mreB5* gene was amplified from the genomic DNA of *S. citri*. The amplified product was cloned into pHis17 vector (for the sequence refer to Addgene catalog # 78202) between *NdeI* and *BamHI* restriction site by restriction free cloning method (39). This resulted in a C-terminal hexa-histidine tagged construct with “GSHHHHHH” added after the last residue. Different single/double point mutants as well as the deletion construct were generated by restriction free cloning using suitable primers. The list of clones and primers are in Table S4.

### Expression and purification of ScMreB5

Plasmid containing the gene of interest was transformed into *E. coli* BL21-AI cells. Cultures were grown in LB medium supplemented with ampicillin (final concentration 100 µg/mL) at 37 °C until an O.D._600_ of 0.8-1.0 was reached. The culture was induced with 0.05% arabinose (final concentration) and was further grown at 20 °C for 12 hours post induction. The same procedure was followed for expression of all the mutants. The culture was spun at 6,000 xg and the cell pellets flash frozen and stored at − 80 °C until further use.

For purification, cell pellet from a 2-litre culture was thawed and cells were homogenized in Lysis Buffer (50 mM Tris pH 8, 200 mM NaCl and 10 % glycerol) and sonicated. The lysate was centrifuged at 44,082 xg for 45 mins, 4 °C. Supernatant was loaded on 5 mL Ni-NTA column (HisTrap, GE Healthcare) pre-equilibrated with buffer A (50 mM Tris pH 8, 200 mM NaCl). Hexa-histidine tag present at the C-terminus of the protein facilitated binding to the Ni-NTA column. Bound protein was eluted using a step gradient of 5%,10%, 20%, 50% and 100% of Buffer B (50 mM Tris pH 8, 200 mM NaCl and 500 mM Imidazole) with Buffer A. Fractions containing purest protein were identified on a 12% SDS-PAGE gel. 1 mM ADP and 1 mM MgCl_2_ (final concentrations) were added to those fractions to minimize protein precipitation. The protein was concentrated and imidazole was removed in buffer exchange while concentrating. Finally, the protein was obtained in the buffer composition 50 mM Tris pH 8, 50 mM NaCl, 1 mM ADP and 1 mM MgCl_2_. Protein was flash frozen in aliquots and stored in –80°C until further use. This protein was used for thermal shift assay.

After optimizing the purification protocol on the basis of thermal shift assay, purification with the optimized conditions were performed similarly as described above, with the following changes. In Lysis Buffer, Buffer A and Buffer B, 200 mM NaCl was replaced with 300 mM KCl. Dialysis was performed post Ni-NTA elution for the fractions containing the purified protein against 50 mM Tris pH 8, 300 mM KCl. Also, ADP and MgCl_2_ were not added at any stage during purification. Analytical size exclusion was performed using Superdex75 or 200 (GE Life Sciences). All experiments other than thermal shift assay and crystallization of ADP-bound ScMreB5 were performed using the protein purified without extra addition of ADP and MgCl_2_.

### Crystallization of ScMreB5-ADP

Approximately 1000 conditions were screened from the commercially available crystallization screens (Molecular Dimensions, Hampton Research) using a drop size of 100 nL of protein (5 mg/mL) and 100 nL crystallization condition. Initial hits obtained were further optimized. Well diffracting, needle shaped crystals were obtained in the condition containing 5 mg/mL protein (in 50 mM Tris pH 8, 50 mM NaCl, 1 mM ADP and 1 mM MgCl_2_ buffer at the end of purification steps) crystallized with 2 mM ADP and 2 mM MgCl_2_ in a condition containing 0.15 M Na-K phosphate, 16% PEG 3350, pH 7.8 by hanging drop method at 1:1 ratio. Crystals were frozen in 20% glycerol as the cryoprotectant contained in the parent condition. Data collection and refinement statistics for ScMreB5–ADP are tabulated in Table S1.

The identity of the potassium ion bound to ScMreB5 in the crystal was established using X-Ray fluorescence scanning. ScMreB5–AMP-PNP was crystallized with 2 mM AMP-PNP and 2 mM MgCl_2_ in the buffer containing 0.15 M Na phosphate, 16% PEG 3350, at pH 7.8 by sitting drop method. To wash out any remnants of potassium ions in the cryoprotectant solution, crystals were picked up and successively transferred into three different drops of 20% glycerol cryoprotectant contained in the parent condition before freezing. X-Ray fluorescence scanning was performed at the beamline I-04 Diamond Light Source, UK.

### Structure determination of ScMreB5-ADP

Data was collected at the home source, Rigaku MicroMax-007 HF. Crystal diffracted till 2.3 Å. Data reduction was performed using IMOSFLM (40), scaling using AIMLESS (41) followed by molecular replacement using PHASER (42) accessed through the CCP4 package (43). Refinement was carried out using PHENIX package (44) and model building was done using *Coot* (45). ScMreB5–ADP bound structure (PDB ID 7BVZ) was solved using CcMreB (PDB ID 4CZI) as model for molecular replacement. Composite omit maps for confirming the ligand densities were calculated using PHASER (42).

### Thermal Shift Assay

For increasing the stability of the protein and optimizing purification, thermal shift assay was performed (46). A final concentration of 2.6 µM of protein (ultracentrifuged at 100,000 g for 25 mins, 4 °C) in a 25 µL reaction was used in this assay. 2 µL of 50X SYPRO Orange (Sigma-Aldrich, S5692) was used as a fluorophore. The reaction was carried out in 96 well PCR plates (Bio-Rad) that was sealed with a microseal (Bio-Rad) after addition of all the components. The plate was spun for 30 secs at 4°C before taking the readings. Bio-Rad CFX96 Real-Time System was used for measuring the T_m_ of the protein by monitoring the change in fluorescence of SYPRO Orange as the protein unfolds. The plate was first incubated in the machine at 4°C for 10 mins. Subsequently, readings were taken from a temperature range of 4 - 90°C with a rise of 0.4°C every 20 secs. Data for the change in SYPRO fluorescence emission as it binds to the hydrophobic pockets of the protein was collected using the FRET channel (excitation at 470 nm and emission at 569 nm). The raw data of the first derivative of the melting curve (-(dF)/dT) was plotted with respect to temperature. GraphPad Prism version 5.00 for Windows, GraphPad Software, San Diego California USA, www.graphpad.com, was used for plotting the graphs.

### Phosphate release assay

The release of inorganic phosphate during ATP hydrolysis was measured by malachite green assay (47). Protein was pre-spun at 22000 xg at 4 °C for 20 mins. Protein was added at a final concentration of 10 µM to the master mix of buffer containing ATP and MgCl_2_ to achieve final concentrations of 1 mM ATP, 1mM MgCl_2_ in Buffer A (300 mM KCl, 50 mM Tris pH 8), and mixed. To stop the reaction for 0 time point reading, 20 µL reaction was immediately mixed with 5 µL of 0.5 M EDTA in a 96 well plate. Rest of the master mix was incubated at 25 °C for 60 mins. After 60 mins, the reaction was stopped with 0.5 M EDTA in the same manner. Simultaneously, phosphate standards were freshly diluted from 400 µM NaH_2_PO_4_ (Sigma-Aldrich, S0751). To measure the amount of phosphate release, malachite green solution was freshly prepared using 800 µL of 3.5 mM Malachite green (Sigma-Aldrich, 38800) dissolved in 3N H_2_SO_4_, 16 µL of 11% Tween 20 (Sigma-Aldrich, P-1379) and 200 µL of 7.5% w/v ammonium molybdate (Sigma-Aldrich, 277908). 50 µL of malachite green solution was added to the stopped reactions and phosphate standards. Absorbance of malachite green was measured in Varioskan Flash (4.00.53) after 5 – 8 mins post addition at 630 nm wavelength.

For calculating the k_obs_ (min^-1^), firstly, the slope was calculated from the phosphate standards. The absorbance of protein containing reaction was calculated by subtracting the blank reaction absorbance (without protein). The amount of phosphate release (in µM) after 60 mins was calculated by dividing the subtracted absorbance with the slope. To calculate the phosphate release per min (k_obs_), the amount of phosphate release (in µM) from protein was divided by 60 mins (µM min^-1^) and then by 10 µM (min^-1^).

GraphPad Prism version 5.00 for Windows, GraphPad Software, San Diego California USA, www.graphpad.com, was used for statistical analysis and plotting the graphs. Statistical significance was estimated by unpaired t-test, two tailed. The data in the graph is expressed as the mean ± standard error in mean (SEM).

### Liposome preparation

All the lipids used in the experiments were purchased from Avanti Polar Lipids, 1,2-dioleoyl-sn-glycero-3-phosphocholine (DOPC, 850375C), 1,2-dioleoyl-sn-glycero-3-phospho-(1′-rac-glycerol) (DOPG, 840475C), porcine brain sphingomyelin (SM, 860062C), *E.coli* Cardiolipin (841199C). For performing pelleting assay with liposomes mimicking *S.citri* lipid composition, protocol described in (23) was followed.

For other assays, a stock concentration of 2 mM lipids in chloroform was made from only DOPG, only DOPC and varying percentage ratios of DOPC:DOPG lipids. Chloroform solution of lipids was aliquoted in a clean test tube and dried. Dried lipids were resuspended in Buffer A (50 mM Tris pH 8, 300 mM KCl) and 1 mM MgCl_2_. This lipid solution was extruded through 100 nm polycarbonate membrane (Avanti Polar Lipids) to get liposomes of 100 nm range. These liposomes were further used in the charge specificity and nucleotide dependent liposome pelleting assays.

### Liposome pelleting assay

Protein was spun at 22000 xg at 4°C for 20 mins to remove any precipitation, if any. From the supernatant, 2 µM (final concentration) protein was added to the reaction mixture of 100 µL containing Buffer A (300 mM KCl, 50 mM Tris pH 8), 1 mM MgCl_2_ and liposomes. This mixture was further incubated at 25°C for 15 mins and spun at 100,000 xg for 25 mins at 25°C. Supernatant was removed and pellet was resuspended in 50 µL Buffer A. Supernatant and pellet were mixed with 2X Laemmli Buffer and equal amounts of both were loaded onto the 12% SDS-PAGE gel. This protocol was followed for the following assays: i) to compare the binding of ScMreB5^WT^ and mutants with liposome composition resembling *Spiroplasma* membrane composition at 1 mM liposomes, ii) to determine charge-based binding specificity of ScMreB5^WT^ and mutants with 1 mM DOPG and 1 mM DOPC liposomes, iii) for determining the binding affinity of ScMreB5^WT^ with increasing concentrations of liposomes (0-1 mM).

The intensity analysis of protein band was performed in Image J 1.52n (48). For calculating the fraction of protein in the pellet, band intensity in the pellet fraction was divided by the sum of band intensities in pellet and supernatant. The data in the graph is expressed as the mean ± standard error in mean (SEM). Statistical significance was estimated by unpaired t-test, two-tailed. GraphPad Prism version 5.00 for Windows, GraphPad Software, San Diego California USA, www.graphpad.com, was used for plotting the graphs.

### Sequence and structure analyses

For domain-wise comparison of ScMreB5-ADP and ScMreB5-AMPPNP with CcMreB, UCSF Chimera version 1.13.1 (49) was used. Each subdomain of ScMreB5 was individually superposed on CcMreB subdomains using the Match Maker option in Chimera in the default settings. The RMSD values obtained for pruned C-alpha atom pairs (generated by iteratively pruning C-alpha atom pairs until each atom pair were within a 2 Å distance cut-off) in each subdomain were tabulated. Figures of crystal structures were prepared using UCSF Chimera version 1.13.1 (49). Sequence alignment of MreBs and actin (Fig. S2) was obtained using Clustal Omega (50) and the figure generated using ESPript 3.0 (51).

For ScMreB5 C-terminus sequence conservation analysis, protein sequences of MreB5s of *Spiroplasma* species listed in Harne et al, 2020 (25) were included for the sequence alignment. All the sequences were aligned using Clustal Omega (50) in default settings. The alignment generated was analyzed and edited in JalView (52). The C-terminus alignment conservation figure (Fig. 5B) was generated using WebLogo (53).

For generating double protofilament assembly model of ScMreB5-AMPPNP, coordinates of the double protofilament assembly of CcMreB (PDB ID 4CZJ) was generated by displaying and saving the coordinates of the symmetry mates using *Coot* (45). Each subdomain of ScMreB5–AMPPNP was saved as separate PDB files through UCSF Chimera version 1.13.1 (49). Each subdomain (IA, IB, IIA, IIB) of ScMreB5–AMPPNP was superposed on one of the protofilament monomers of the CcMreB filament using Match Maker option in Chimera in the default settings. Similarly, ScMreB5 subdomains were superposed on the other two monomers of the same protofilament of CcMreB double protofilament assembly. This generated a single protofilament of ScMreB5-AMPNP in a double protofilament model. Second protofilament of ScMreB5 was also generated by repeating the same steps. This resulted in a double protofilament model of ScMreB5-AMPPNP. Electrostatic potential surface of this model was generated using “Electrostatic surface coloring” option in Chimera with the default settings.

### Cryo-electron microscopy

For visualizing the filaments of ScMreB5^WT^ and the ATPase mutant ScMreB5^E134A^, electron cryomicroscopy was carried out. Quantifoil Au 1.2/1.3 grids that were glow discharged for 90 secs were used. Protein was ultracentrifuged at 100,000 g for 25 mins, 4 °C. 5 mM nucleotide (AMP-PNP, ADP or ATP) and 5 mM MgCl_2_ (final concentrations) were added to a final concentration of 50 µM protein and incubated at 25 °C for 10-15 mins. 3 µL of the sample was put on the grid and incubated for 5-10 s before blotting for 3 secs followed by plunge freezing into liquid ethane for vitrification using a FEI Vitrobot. For image acquisition, grids were mounted on Triton-Krios 300 KeV electron microscope with Falcon-3 direct electron detector and images were taken at a magnification of 59,000X. Images of the filaments were generated on ImageJ 1.52n (48).

### Yeast strains and growth conditions

GFP-ScMreB5^WT^ and GFP-ScMreB5^E134A^ were expressed from the thiamine repressible medium-strength *nmt41/42* promoter (54). *pREP41-GFP-ScMreB^WT^* and *pREP41-GFP-ScMreB^E134A^* expression vectors were transformed in *Schizosaccharomyces pombe* strain MBY192 (*h^−^ leu1-32 ura4-D18*; lab collection) using Li-acetate method (55). Yeast cells were grown on Edinburgh minimal medium (EMM) with the addition of specific supplements (histidine, uracil, adenine). Cultures were grown in the absence of thiamine as indicated below in EMM to allow expression of the protein.

### Light Microscopy

*S. pombe* strains carrying GFP-ScMreB5^WT^, GFP-ScMreB5^E134A^ were grown in EMM broth for 24 – 32 hours at 36°C in the absence of thiamine for the expression of protein and imaging. For counting of cells and comparison of number of cells with polymer bundles, cultures were grown for 36 – 48 hours. At least 1000 cells were counted for each sample and the experiment was repeated at least thrice. Cultures were intermittently diluted into fresh medium to maintain exponential growth. For live cell microscopy, 1 mL of cells after 24 – 32 hours of growth in the absence of thiamine were pelleted down at 855 xg for 2 minutes and reconstituted in 50 - 100 µL of fresh medium. 2 µL of cell suspension was then mounted on EMM agarose slides, a coverslip placed and sealed with VALAP. Further images were collected in Z-step of 0.2 µm or 0.5 µm at a fixed 3 minutes interval of time for 6 – 12 hours by using an epifluorescence microscope (DeltaVision^TM^ Elite) equipped with 100X 1.4 NA oil-immersion objective. Images were acquired using an interline CCD camera Photometrics® CoolSnap HQ2^TM^. GFP-ScMreB5 was imaged using excitation and emission filters of 475/28 nm and 525/48 nm, respectively. UltimateFocus^TM^ was used to maintain the cells in focus during the entire duration of imaging. Deconvolution was performed using the SoftWorx^TM^ software. 3D-SIM was performed using DeltaVision^TM^ OMX-SR Blaze with cells mounted on an agarose pad as described above. Raw images were acquired using a 60X NA 1.42 oil-immersion objective lens and a PCO Edge 4.2 sCMOS camera and reconstructed using the SI reconstruction module of SoftWorx^TM^ software. All images were processed by using Fiji software (v 2.0.0-rc-69/1.52p) (56). GraphPad Prism version 5.00 for Windows, GraphPad Software, San Diego California USA, www.graphpad.com, was used for plotting the graph.

## Acknowledgments

This work was initiated with the support from Department of Science and Technology INSPIRE Faculty Fellowship (IFA12/LSBM-52), Innovative Young Biotechnologist Award (BT/07/IYBA/2013) to PG. The work is currently funded by Department of Biotechnology (DBT) Membrane Structural Biology Program grant (BT/PR28833/BRB/10/1705/2018) and IISER Pune to PG, and SERB (EMR/2016/000487), DBT (BT/PR15183/BRB/10/1443/2015) and intramural core funding from Department of Atomic Energy (DAE) to RS. Macromolecular crystallography facility at IISER Pune and synchrotron facilities at ESRF, Grenoble, Diamond Light Source (MX22637) and EMBL-ESRF and DBT India for facilitating data collection at ID29, ESRF and the National Electron Cryomicroscopy Facility, Bangalore Life Sciences Cluster, India (K.R.Vinoth Kumar and Mamta Bangera) are acknowledged. Staff members at the Centre of Interdisciplinary Sciences, NISER for routine maintenance of OMX-SR Super-resolution imaging facility and Ajay Kumar Sharma for initial help with 3D-SIM imaging are acknowledged. We also acknowledge fellowships from IISER Pune and Infosys Foundation (VP), NM (DAE), INSPIRE (SRB).

## Author contributions

VP designed and performed all experiments other than those mentioned below, analyzed the data and wrote the manuscript; NM performed the yeast microscopy experiments; SRB standardized the initial purification of ScMreB5 and crystallized the ScMreB5-ADP complex; RS designed, analyzed and supervised the yeast microscopy experiments; NM & RS wrote the sections pertaining to yeast experiments; PG designed, conceptualized, supervised the study and wrote the manuscript. All authors reviewed and provided inputs on the manuscript.

