## Supplemental Tables and Figures for "Filament dynamics driven by ATP hydrolysis modulates membrane binding of the bacterial actin MreB"

**This PDF file includes:**

Supplementary Figures S1 to S4
Supplementary Tables S1 to S4
Supplementary Movie Legends S1 to S5

**Supplementary figures**

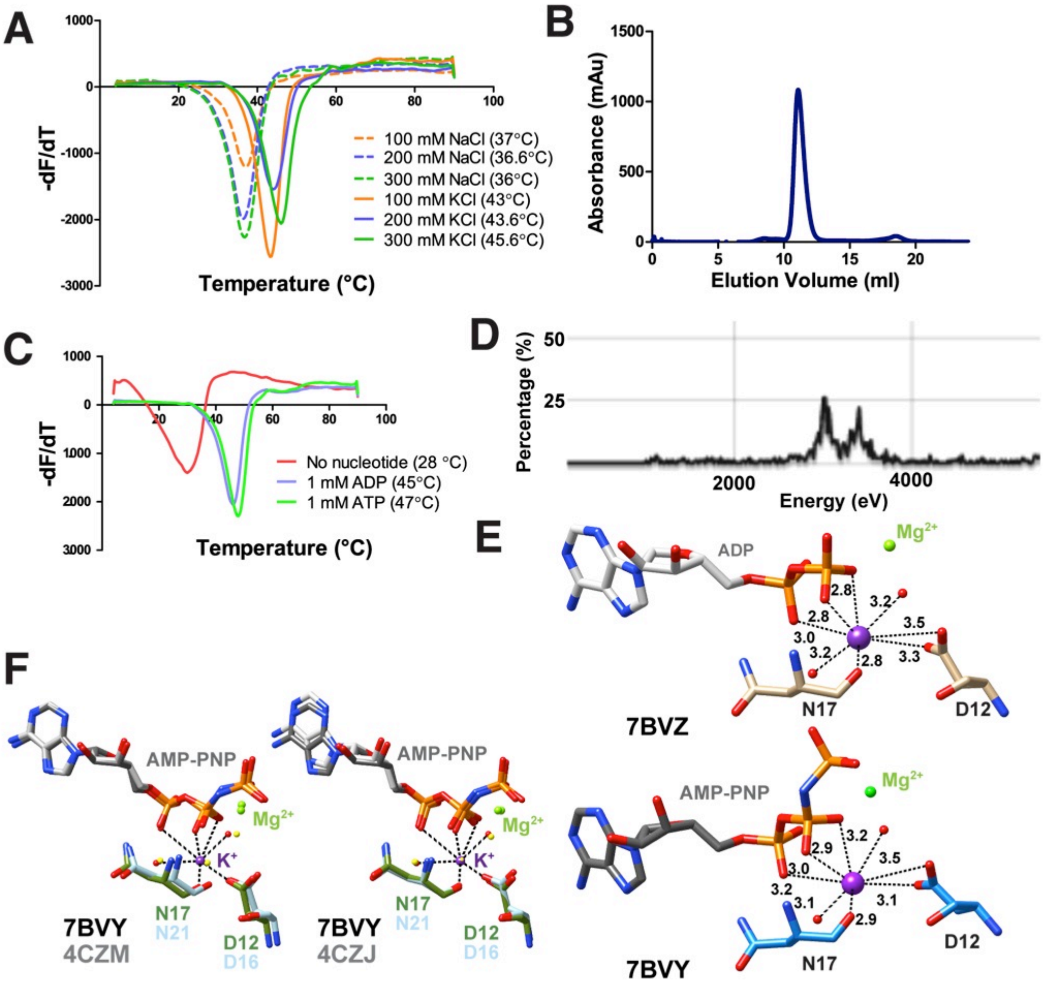


**Supplementary Figure 1. ScMreB5 is stabilized by KCl and nucleotides.**

**A.** Melting curve for ScMreB5 showing $T_m$ for varying concentrations (orange - 100 mM; blue - 200 mM and green - 300 mM) of NaCl (dotted line) and KCl (solid line).

**B.** Analytical size exclusion chromatography using Superdex75 (GE LifeSciences) for ScMreB5 in Buffer A (300 mM KCl, 50 mM Tris, pH 8.0) shows a single peak corresponding to monomeric ScMreB5.

**C.** Melting curve for ScMreB5 showing $T_m$ of ScMreB5 without any nucleotide (red), 1 mM ADP (blue) and 1 mM ATP (green).

**D.** X-Ray fluorescence scan for ScMreB5–AMP-PNP crystals, showing the signature for potassium.

**E.** Coordination sphere of potassium in ScMreB5–ADP (PDB ID: 7BVZ; top) and ScMreB5–AMPPNP (PDB ID: 7BVY; bottom).

**F.** Asp12 and Asn 17 of ScMreB5 at the potassium binding site are compared with the corresponding residues present in monomeric (PDB ID 4CZM) and double protofilament CcMreB (PDB ID 4CZJ) by superposing IIA subdomain of CcMreBs onto IIA domain of ScMreB5–AMPPNP structure (single protofilament conformation). Presence of water molecule in the CcMreB at the potassium equivalent position can be observed. The residues of ScMreB5 are coloured domain wise and of CcMreB in light blue. The water molecules for ScMreB5 are coloured in red and for CcMreBs in yellow.

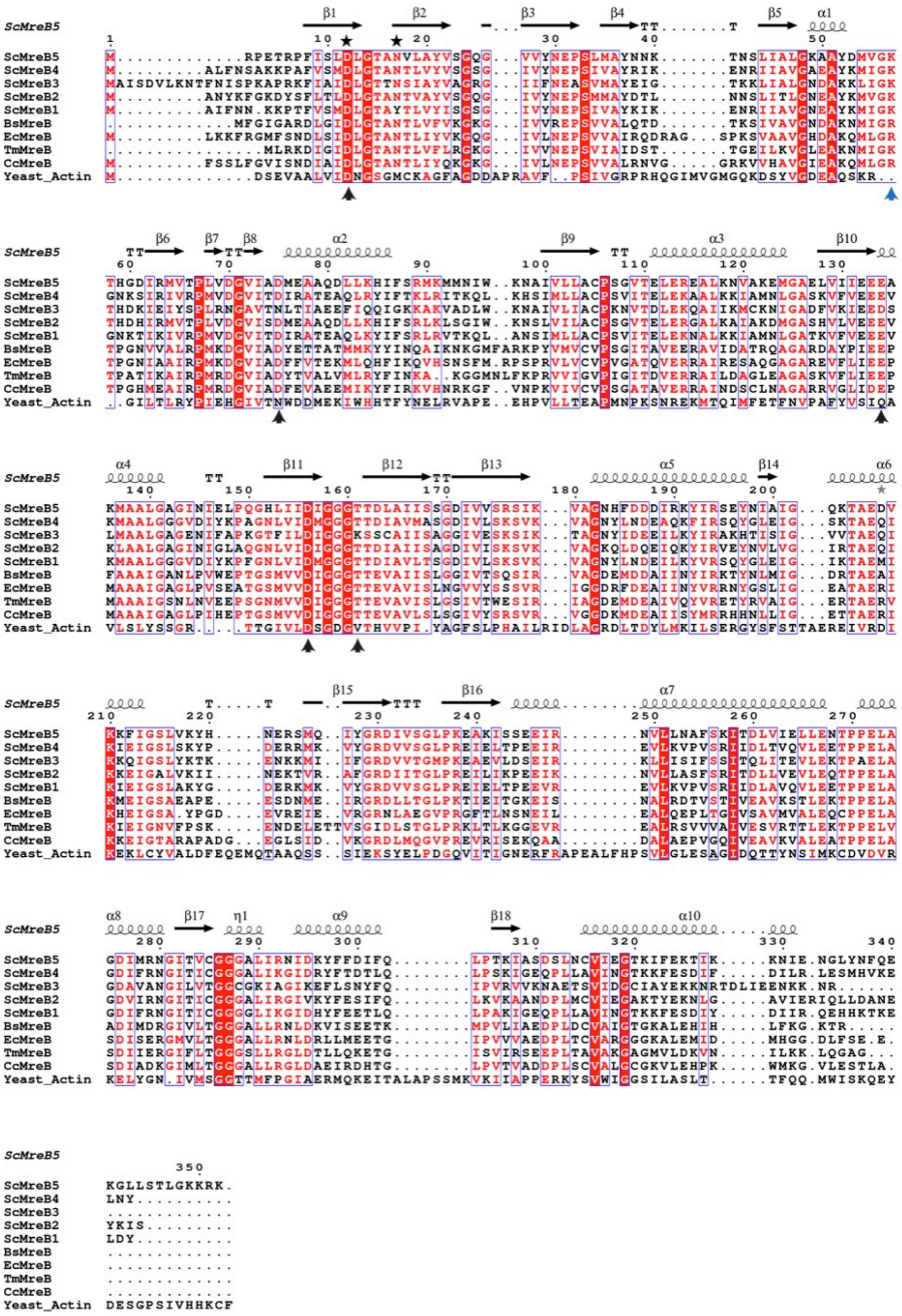

**Supplementary Figure 2: Residues at nucleotide binding pocket are well conserved in ScMreB5**

Sequence alignment of ScMreB5 (ScMreB5) with other *S. citri* MreBs (ScMreB1, ScMreB2, ScMreB3 and ScMreB4), *C. crecentus* MreB (CcMreB), *T. maritima* MreB (TmMreB), *E. coli* MreB (EcMreB), *B. subtilis* MreB (BsMreB) and Yeast actin. Residues mutated with respective roles in ATP hydrolysis (black arrowheads), $K^+$ coordination (black star) and polymerization interface (blue arrowheads) are marked.

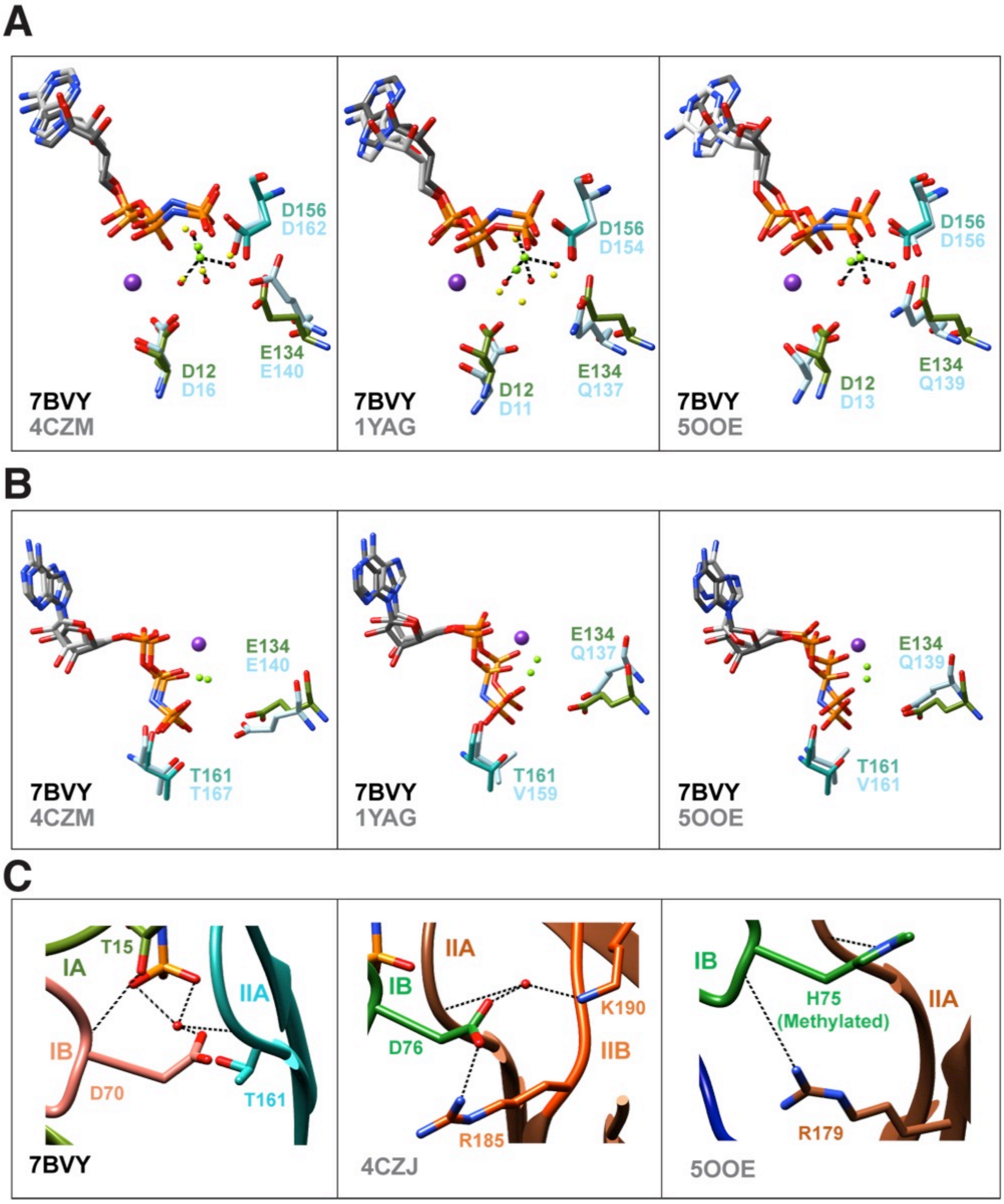


**Supplementary Figure 3. Comparison of active site residues of ScMreB5 with CcMreB and actin.**

Active site residues of ScMreB5 are compared with the monomeric CcMreB (PDB ID 4CZM), double protofilament CcMreB (PDB ID 4CZJ) monomeric yeast actin (PDB ID 1YAG) and actin filament (PDB ID 5OOE) by superposing IIA domain of each structure onto IIA domain of ScMreB5-AMPPNP structure (PDB ID 7BVY; single protofilament conformation).

**A.** Superposed active site residues holding the $Mg^{2+}$ coordination sphere. Asp12, Glu134 and Asp156 residues of ScMreB5 are compared with the corresponding residues in CcMreB and actin structures.

**B.** Superposed active site residues at the vicinity of the catalytic water. Glu134 and Thr161 conformations in ScMreB5 are compared with the corresponding residues in CcMreB and actin structures. ScMreB5-AMP-PNP residues are colored subdomain-wise for IA (green), IB (pink) and IIA (sea green). For the CcMreB and actin structures, corresponding residues in subdomains are colored light blue.

**C.** Interacting interface for Asp70 of ScMreB5-AMP-PNP (7BVY) is shown with respect to corresponding residues present in CcMreB double protofilament (PDB ID 4CZJ) and actin filament (PDB ID 5OOE).

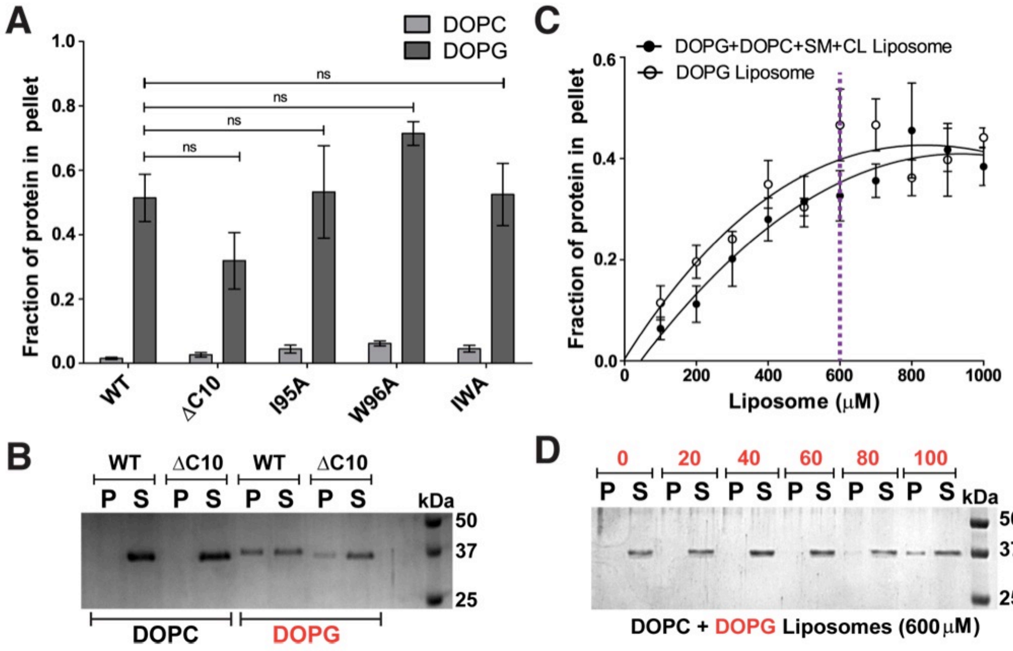


**Supplementary Figure 4. Binding of ScMreB5 to DOPG liposome is specific and concentration dependent.**

**A.** Plot showing relative intensities of the fraction of protein in the pellet corresponding to ScMreB5$^{WT}$ and mutant constructs in the SDS-PAGE gels from 5 independent experiments. The binding is specifically observed for liposome composed of the anionic lipid DOPG for the ScMreB5$^{WT}$ as well as the mutants. Negligible binding is seen for both ScMreB5$^{WT}$ as well as the mutants for the liposome made from neutral lipid DOPC. The error bar denotes the mean with standard error of the mean.

**B.** A representative 12% SDS-PAGE gel of liposome pelleting assay with C-terminal deletion mutant (ΔC10) shows binding with the charged liposome, DOPG. P and S represent the pellet and supernatant fractions of the protein.

Concentration of DOPG and DOPC liposomes used in the assay is 1 mM and protein is 2 μM.

**C.** Liposome binding curves showing the increase in the fraction of ScMreB5 WT in the pellet (liposome bound fraction) at 2 μM protein concentration with increasing concentration of the liposomes mimicking *Spiroplasma* lipid composition and DOPG liposomes. 600 μM liposome concentration was maintained as a constant for further experiments performed by varying the liposome composition.

**D.** A representative 12% SDS-PAGE gel of liposome pelleting assay for showing the binding specificity of $ScMreB5^{WT}$ by varying the DOPC and DOPG ratios at 600 μM liposome concentration. Protein in the pellet is observed at the higher DOPG percentages (labeled in red on top). P and S represent the pellet and supernatant fractions of the protein.

### Supplementary Tables

#### Supplementary Table 1: Data collection and refinement statistics of *S. citri* MreB5 bound to ADP (PDB ID: 7BVZ)

| ScMreB5–ADP (PDB ID: 7BVZ) | |
|---|---|
| | ***Data Collection statistics*** |
| Collected at | Rigaku Micromax-007 |
| Wavelength (Å) | 1.5418 |
| Space Group | P1 |
| a, b, c (Å) | 37.9, 41.1, 56.3 |
| α, β, γ (˚) | 82.3, 74, 80.4 |
| Resolution (Å)* | 40.4-2.3 (2.38 – 2.3) |
| Number of unique reflections* | 12584 (1233) |
| $R_{merge}$ (%)* | 0.12 (0.63) |
| $R_{pim}$ (%)* | 0.11 (0.52) |
| $CC_{half}$* | 0.98 (0.48) |
| Mean I/ ʃI | 6.1 (1.5) |
| Completeness (%)* | 87.9 (86.2) |
| Redundancy* | 2.1 (2.1) |
| ***Refinement statistics*** | |
| Resolution (Å) | 36.14-2.3 |
| Number of unique reflections (test set) | 12580(657) |
| $R_{work}$ / $R_{free}$ (%) | 22.5/27.2 |
| Average B-factor (Å$^2$) | 23.1 |
| Wilson B-factor (Å$^2$) | 19.22 |
| *RMS deviations* | 0.002 |
| Bond lengths (Å) | 0.51 |
| Bond angles (˚) | |
| *Ramachandran map statistics* | |
| Favored (%) | 97.85 |
| Allowed (%) | 7 |
| Outliers (%) | 0 |

* *Values in parentheses denote the last resolution shell.*

#### Supplementary Table 2: Subdomain wise RMSD of ScMreB5 structures compared with CcMreB (PDB ID: 4CZI)

| Subdomains | ScMreB5-AMPPNP | Pruned atoms | ScMreB5-ADP | Pruned atoms |
|---|---|---|---|---|
| IA | 1.07 | 82 | 1.05 | 85 |
| IB | 0.97 | 26 | 1.1 | 31 |
| IIA | 0.99 | 96 | 1.01 | 98 |
| IIB | 0.62 | 65 | 0.69 | 64 |

### Supplementary Table 3: $k_{obs}$ values of ScMreB5$^{WT}$ and mutants

| Protein | Activity, $k_{obs}$ ($min^{-1}$) |
|---|---|
| ScMreB5$^{WT}$ | 0.15 ± 0.007 (Active) |
| ScMreB5$^{D12A}$ | 0.01 ± 0.008 (Inactive) |
| ScMreB5$^{D70A}$ | 0.02 ± 0.002 (Inactive) |
| ScMreB5$^{E134A}$ | 0.05 ± 0.005 (Inactive) |
| ScMreB5$^{D156A}$ | 0.08 ± 0.014 (Inactive) |
| ScMreB5$^{T161A}$ | 0.0005 ± 0.001(Inactive) |
| ScMreB5$^{K57A}$ | 0.1 ± 0.01 (Partially active) |

### Supplementary Table 4: List of primers

| Primer name | Sequence ('5 → 3') | Clones generated using the primers (Vector-construct name) |
|---|---|---|
| ScM5-f | CTTTAAGAAGGAGATATACATATGAGACCAG AAACTAGACCATTTATTTC | pHis17-ScMreB5$^{WT}$ |
| ScM5H6-r | GATGATGATGATGATGGGATCCTTTTCTTTTT TTACCTAATGTTGATAATAATCC | |
| ScM5W-f | GAATGAAAATGATGAACATTTGGAAGAATGC TATTG | |
| ScM5-k57a-f | CTATGATATGGTAGGAGCAACACACGGAGA TATTAG | pHis17-ScMreB5$^{K57A}$ |
| ScM5-D156-f | GGTCATTTAATCATTGCTATCGGTGGAGGAA CAAC | pHis17-ScMreB5$^{D156A}$ |
| ScM5-E134-f | GTTATCATTGAAGAAGCGGCTAAAATGGCCG | pHis17-ScMreB5$^{E134A}$ pRep-ScMreB5$^{E134A}$-NGFP |
| D12A-f | CCAGAAACTAGACCATTTATTTCTCTTGCGTT AGGAACTGCTAATG | pHis17-ScMreB5$^{D12A}$ |
| ScM5-D70-f | GGTAACACCATTAGTAGCGGGAGTTATCGC AGACATGGAAGCTGCAC | pHis17-ScMreB5$^{D70A}$ |
| ScM5-T161A-f | GGTGGAGGAGCGACTGATTTAGCTATTATTT CATCAGGTG | pHis17-ScMreB5$^{T161A}$ |
| M5-I95A | CAAGAATGAAAATGATGAACGCGTGGAAGA ATGCTATTGTATTATTAGC | pHis17-ScMreB5$^{I95A}$ |
| M5-W96A | CAAGAATGAAAATGATGAACATTGCGAAGAA TGCTATTGTATTATTAGC | pHis17-ScMreB5$^{W96A}$ |
| M5-IWA | CAAGAATGAAAATGATGAACGCCGCGAAGA ATGCTATTGTATTATTAGC | pHis17-ScMreB5$^{IWA}$ |
| ScM5Ct10 del-r | GCTTTTAATGATGATGATGATGATGGGATCC TTTTTCTTGAAAATTATATAAACC | pHis17-ScMreB5$^{\Delta C10}$ |
| pREP_ScM5-f | GGCATGGATGAACTATACAAACATATGATGA GACCAGAAACTAGACCATTTATTTC | pRep-ScMreB5-NGFP |
| pREP_ScM5-r | GGCAAGGGAGACATTCCTTTTACCCGGGGA TCCTTATTTTCTTTTTTTACCTAATGTTG | |

**Supplementary Movies**

**Movie S1** (Corresponding frames shown in Fig. 3B).

**360° Volume-rendering 3D-SIM images of ScMreB5$^{WT}$ and ScMreB5$^{E134A}$ filaments.**

360° volume rendering of GFP-ScMreB5$^{WT}$ and ScMreB5$^{E134A}$ filaments in fission yeast. 3D-SIM reconstruction of the images was carried out using SoftWorx$^{TM}$ software. 3D volume data was constructed using FIJI software. Maximum Intensity projection image is shown in Fig. 3B. Scale bar represents 5 µm.

**Movie S2** (Corresponding frames shown in Fig. 3D)

**Polymerization of ScMreB5$^{WT}$ and ScMreB5$^{E134A}$ in *S. pombe* cells.**

Time-lapse series showing the polymerization of GFP-ScMreB5$^{WT}$ and GFP-ScMreB5$^{E134A}$ in *S. pombe* cells observed by epi-fluorescence microscopy. Panels shown in Fig. 3D (WT and E134A) were obtained from this time-series. Images were deconvolved using SoftWorx$^{TM}$ software and are maximum intensity projections from Z-stacks of 0.5 µm and acquired 3 minutes interval. Scale bar represents 5 µm.

**Movie S3** (Corresponding frames shown in Fig 3E)

**ATP hydrolysis mutant ScMreB5$^{E134A}$ shows defects in bundling of filaments compared to ScMreB5$^{WT}$ filaments.**

Time-lapse microscopy showing bundling of GFP-ScMreB5$^{WT}$ filaments. GFP-ScMreB5$^{WT}$ filaments make lateral contacts and bundle. Septation is often seen to bring filaments together and act as a trigger for bundling as well. GFP-ScMreB5$^{WT}$ filaments fail to bundle. Although septation is often seen to bring filaments together, it fails to induce bundling of GFP-ScMreB5$^{E134A}$ filaments. Panels shown in Fig 3E were obtained from this time-series.

Images were deconvolved using SoftWorx$^{TM}$ software and are maximum intensity projections from Z-stacks of 0.2 μm and acquired 3 minutes intervals. Scale bar represents 5 μm.

**Movie S4** (Corresponding frames shown in Fig 3F)

**Disassembly of ScMreB5$^{WT}$ filaments.**

Time-lapse series showing disassembly of GFP-ScMreB5$^{WT}$ filaments in *S. pombe* cells. Panels shown in Fig 3F were obtained from this time-series. GFP-ScMreB5$^{WT}$ filaments are seen to disassemble or depolymerize, probably owing to dilution of protein concentration immediately upon cell division. Cells were observed by epi-fluorescence microscopy and images were deconvolved using SoftWorx$^{TM}$ software. Images shown are maximum intensity projections from Z-stacks of 0.5 μm and acquired 3 minutes interval. Scale bar represents 5 μm.

**Movie S5** (Corresponding frames shown in Fig 3G)

**Fragmentation and annealing of ScMreB5$^{WT}$ filaments.**

Time-lapse series showing fragmentation and annealing of GFP-ScMreB5$^{WT}$ filaments in *S. pombe* cells. Panels shown in Fig 3G were obtained from this time-series. GFP-ScMreB5$^{WT}$ filaments are seen to fragment and re-anneal. Cells were observed by epi-fluorescence microscopy and images were deconvolved using SoftWorx$^{TM}$ software. Images shown are maximum intensity projections from Z-stacks of 0.5 μm and acquired 3 minutes interval. Scale bar represents 5 μm.
